# Effects of Neonicotinoid Seed Treatments on Soil Microbial Gene Expression Vary with Time in an Agricultural Ecosystem

**DOI:** 10.1101/2022.01.20.477174

**Authors:** Mona Parizadeh, Benjamin Mimee, Steven W. Kembel

## Abstract

Neonicotinoids, a class of systemic insecticides, have been widely used for decades against various insect pests. Past studies have reported non-target effects of neonicotinoids on some beneficial macro- and micro-organisms. Given the crucial role that the soil microbiota plays in sustaining soil fertility, it is critical to understand how microbial taxonomic composition and gene expression respond to neonicotinoid exposure. To date, few studies have focused on this question, and these studies have evaluated the shifts in soil microbial taxonomic composition or used soil biochemical analyses to assess the changes in microbial functions. In this study, we have applied a metatranscriptomic approach to quantify the variability in soil microbial gene expression in a two-year soybean/corn crop rotation in Quebec, Canada. We identified weak and temporally inconsistent effects of neonicotinoid application on soil microbial gene expression, as well as a strong temporal variation in soil microbial gene expression among months and years. Neonicotinoid seed treatment altered the expression of a small number of microbial genes, including genes associated with heat shock proteins, regulatory functions, metabolic processes and DNA repair. These changes in gene expression varied during the growing season and between years. Overall, the composition of soil microbial expressed genes seems to be more resilient and less affected by neonicotinoid application than soil microbial taxonomic composition. Our study is among the first to document the effects of neonicotinoid seed treatment on microbial gene expression and highlights the strong temporal variability of soil microbial gene expression and its responses to neonicotinoid seed treatments.

**IMPORTANCE:** This work provides the first example of the impacts of neonicotinoid seed treatment on community-wide soil microbial gene expression in an experimental design representing real farming conditions. Neonicotinoid pesticides have attracted a great deal of attention in recent years due to their potential non-target impacts on ecological communities and their functions. Our paper represents the first use of metatranscriptomic sequencing to offer real-time and in-depth insights into the non-target effects of this pesticide on soil microbial gene expression and on potentially beneficial soil microorganisms.

## INTRODUCTION

Soil quality is frequently used as an indicator of environmental health in sustainable agriculture (1). It refers to the capacity of soil to function in order to sustain biological productivity and maintain or improve environmental quality and the health of humans, plants, animals, and other living organisms (2). Soil microbial diversity, composition and functions are important indicators to monitor and evaluate soil quality (3,1). Ecological disturbances caused by environmental stress and perturbations such as pesticide application have been shown to influence microbial community structure and functional diversity (4, 5). To better understand the effects of these disturbances on soil microbiome, it is crucial to study microbial functional activities and gene expression (6). Past studies have reported effects of some pesticides on soil microbial functional activities such as microbial biomass enzyme activities and biochemical reactions, including carbon or nitrogen mineralization, nitrogen fixation, nitrification, and denitrification (7, 5). However, to date, a systematic evaluation of the effects of pesticide application on community-wide soil microbial gene expression is lacking. Here we address this lack of knowledge by measuring the effects of neonicotinoid application and temporal variation on soil microbial gene expression in a soybean-corn agroecosystem in Quebec.

Neonicotinoids are a widely used family of systemic neuro-active insecticides that are chemically similar to nicotine. They were introduced to the world in the late 1980s (8, 9) and today, they are used prophylactically in the form of seed treatments against a variety of insect pests (10, 11, 12). Past studies have shown the non-target effects of these pesticides on beneficial insect pollinators such as honeybees and butterflies, and soil invertebrates such as earthworms (13, 14, 15, 16, 17, 18). Neonicotinoids are supposed to be selectively more toxic to invertebrates because of the fundamental distinctions between their nicotinic acetylcholine receptors (nAChRs) compared to vertebrates (9). However, non-target impacts of these pesticides on the taxonomic composition of soil microbial communities have been documented, including shifts in the abundance of diverse taxa, such as a decrease in bacteria genera involved in nitrification and an increase in bacteria genera related to neonicotinoid biodegradation (19, 20, 21, 22, 23, 24, 25). An increase in the abundance of the genes coding for the cytochrome P450 enzyme family has been reported in response to neonicotinoid exposure, based on soil microbial amplicon and metagenomic sequencing (26, 27). Previous studies have indicated that this family of detoxifying enzymes is also over-expressed in the insects resistant to this pesticide and is involved in neonicotinoid biodegradation (28, 29, 30). Another study has reported that nitrogen-fixing and nitrifying bacteria are very sensitive to neonicotinoids (31). Studies on the effects of neonicotinoids on gene expression in different plant species have shown a variety of responses, including a decrease in the expression of cell wall synthesis-related genes, which may lead to a lower resistance to cell-content feeder insects, and an increase in the expression of (1) photosynthesis-related genes, which may prolong the energy production period, (2) pathogenesis-related genes, and (3) stress tolerance-related genes (for example genes involved in tolerance to drought and cold) (32,33,34,35). However, these changes are not consistent and their mechanisms are unknown (36, 37).

To our knowledge, none of these past studies have quantified community-wide changes in soil microbial gene expression in response to neonicotinoid seed treatment; rather, they have focused on the expression of one or a few genes at a time. Similarly, biochemical studies have shown that neonicotinoids can have non-target impacts on soil microbial functional activities and biochemical processes, such as a decline in soil respiration, nitrification and the activity of nitrite and nitrate reductase enzyme, as well as an inhibition in metabolic processes resulting in a decrease in enzymatic activity (38,39,31,40). But, these studies have focused on one or a few indicators of microbial function. Thus, while there is evidence for changes in individual measures of microbial functional activities, we are not aware of studies that have used transcriptomic or metatranscriptomic approaches to quantify community-wide changes in soil microbial gene expression in response to neonicotinoid seed treatment.

In this study, we used metatranscriptomics to evaluate the effects of neonicotinoid seed treatment on soil microbial gene expression. Metatranscriptomics (also known as RNA-seq) identifies the genes that are actually being expressed in a given environment and can help to better study the active functions and the adaptations of microbial communities to environmental changes and stress (41, 42, 43). In this study, our specific objectives were to (1) characterize soil microbial gene expression, including bacterial and eukaryotic expressed genes, in a two-year soybean/corn crop rotation using metatranscriptomic sequencing, and (2) assess the effects of neonicotinoid seed treatment on soil microbial gene expression in this agroecosystem. We hypothesized that (1) neonicotinoid seed treatment and time affect soil microbial gene expression and (2) the expression of pesticide degradation-related genes increases, while the expression of nitrification-related genes decreases in response to neonicotinoid seed treatment. To address our objectives and hypotheses, we studied soil microbial gene expression using a metatranscriptomic approach in a two-year soybean/corn crop rotation in Quebec, Canada.

## RESULTS

### Soil microbial profiling based on SEED hierarchical microbial functional and RefSeq bacterial and eukaryotic functional categories

We detected an average (mean ± SE) of 4,878 ± 4 SEED hierarchical functional categories (level 4) per sample, 22,902 ± 162 RefSeq bacterial functional categories per sample, and 9,899 ± 206 RefSeq eukaryotic functional categories per sample. The SEED-based hierarchical annotation results indicated that 50.5% of the total relative abundance of microbial expressed genes at level 1 of the SEED hierarchy belonged to the ten most abundant microbial functional categories at this level (Table 1A). The majority of the most abundant level 4 SEED hierarchy functional categories were similar to the ten most abundant bacterial and eukaryotic RefSeq-based functional categories, including genes related to chaperone GroEL, chaperone DnaK, DNA-directed RNA polymerase beta subunit, elongation factor G and elongation factor T (Table 1B and Fig. S1). The ten most abundant functional categories accounted for 21.7%, 10.0% and 18.1% of the total relative abundance of, respectively, SEED hierarchical microbial (level 4), RefSeq bacterial and eukaryotic expressed genes (Table 1B and Fig. S1).

**TABLE 1.** Ten most abundant soil SEED hierarchical functional categories (levels 1-3: A and level 4: B), RefSeq bacterial and eukaryotic functional categories (B) in a two-year soybean/corn crop rotation in L’Acadie, Quebec, Canada.

| Functional Databases | Functional Categories | Relative Abundance (%) |
| --- | --- | --- |
| SEED Hierarchical Profile (Level 1) | Protein biosynthesis | 13.20 |
|  | No hierarchy / NA | 9.67 |
|  | Transcription | 5.44 |
|  | Protein folding | 5.29 |
|  | Clustering-based subsystems | 4.46 |
|  | Central carbohydrate metabolism | 3.56 |
|  | Protein degradation | 2.50 |
|  | Resistance to antibiotics and toxic compounds | 2.38 |
|  | Lysine, threonine, methionine, and cysteine | 2.04 |
|  | Heat shock | 1.93 |
| SEED Hierarchical Profile (Level 2) | No hierarchy / NA | 25.30 |
|  | Protein Metabolism | 21.60 |
|  | Carbohydrates | 9.38 |
|  | Amino Acids and Derivatives | 6.77 |
|  | RNA Metabolism | 6.74 |
|  | Stress Response | 5.33 |
|  | Respiration | 3.83 |
|  | Cofactors, Vitamins, Prosthetic Groups, Pigments | 3.25 |
|  | Virulence, Disease and Defense | 2.54 |
|  | Clustering-based subsystems | 2.15 |
| SEED Hierarchical Profile (Level 3) | No hierarchy / NA | 9.57 |
|  | Ribosome LSU bacterial | 4.60 |
|  | GroEL GroES | 4.42 |
|  | Ribosome SSU bacterial | 3.60 |
|  | RNA polymerase bacterial | 3.02 |
|  | Translation elongation factors bacterial | 1.98 |
|  | Heat shock dnaK gene cluster extended | 1.93 |
|  | Proteolysis in bacteria, ATP-dependent | 1.90 |
|  | Transcription initiation, bacterial sigma factors | 1.63 |
|  | Ton and Tol transport systems | 1.42 |

**B)**
| SEED Hierarchy (Level 4) | Relative Abundance (%) | RefSeq Bacteria | Relative Abundance (%) | RefSeq Eukaryotes | Relative Abundance (%) |
| --- | --- | --- | --- | --- | --- |
| No hierarchy / NA | 9.57 | Molecular chaperone GroEL | 2.59 | Heat shock protein 60, mitochondrial precursor | 4.33 |
| Heat shock protein 60 family chaperone GroEL | 3.80 | DNA-directed RNA polymerase subunit beta | 2.00 | Heat shock protein 78, mitochondrial precursor | 2.12 |
| DNA-directed RNA polymerase beta subunit (EC 2.7.7.6) | 2.57 | Molecular chaperone DnaK | 0.96 | Putative chaperonin GroL | 1.73 |
| Chaperone protein DnaK | 1.21 | ABC transporter ATP-binding protein | 0.91 | Cold-shock DNA-binding domain-containing protein | 1.66 |
| Translation elongation factor Tu | 0.87 | MFS transporter | 0.65 | Chaperonin homolog Hsp-60, mitochondrial | 1.53 |
| RNA polymerase sigma factor RpoD | 0.80 | Elongation factor G | 0.64 | Elongation factor Tu, mitochondrial precursor | 1.53 |
| Translation elongation factor G | 0.76 | Endopeptidase La | 0.60 | Chaperonin Hsp-60 | 1.45 |
| ATP-dependent protease La (EC 3.4.21.53) Type I | 0.75 | ABC transporter substrate-binding protein | 0.57 | Heat shock 60kD protein 1 | 1.40 |
| SSU ribosomal protein S1p | 0.71 | DNA-binding response regulator | 0.56 | Chaperone protein DnaK | 1.30 |
| Cell division protein FtsH (EC 3.4.24.-) | 0.65 | Elongation factor Tu | 0.54 | Chaperonin homolog HSP60, mitochondrial precursor, partial | 1.00 |

### Effects of neonicotinoid seed treatment on the composition and diversity of soil microbial expressed genes

Neonicotinoid seed treatment had no significant effect on the overall composition and diversity of soil microbial expressed genes (based on PERMANOVA and Wilcoxon rank-sum test on Shannon index). However, time (year and month) was an important driver of variation in the composition and diversity of soil microbial expressed genes. Year and month together explained significant variation in gene expression at level 4 of SEED hierarchical functional categories (25.07%), RefSeq bacterial functional categories (21.33%), and RefSeq eukaryotic functional categories (10.90%) (PERMANOVA *P* <0.001, Table 2 and Fig. 1).

**TABLE 2.** Drivers of the soil microbial gene expression variation in response to neonicotinoid seed treatment, time and their interactions in a two-year soybean/corn rotation in l’Acadie, Quebec, Canada (PERMANOVA based on Bray-Curtis dissimilarities).

|  | SEED Hierarchical Gene Expression |  |  | RefSeq Bacterial Gene Expression |  |  | RefSeq Eukaryotic Gene Expression |  |  |
| --- | --- | --- | --- | --- | --- | --- | --- | --- | --- |
| Variables | R<sup>2</sup> (%) | F | Pr(>F) | R<sup>2</sup> (%) | F | Pr(>F) | R<sup>2</sup> (%) | F | Pr(>F) |
| Year/Month | 25.07 | 14.64 | 0.001*** | 21.33 | 10.86 | 0.001*** | 10.90 | 4.65 | 0.001*** |
| Neonicotinoid seed treatment | 1.91 | 1.11 | NS | 2.13 | 1.08 | NS | 1.87 | 0.80 | NS |
| Year/Month : Neonicotinoid seed treatment | NS | NS | NS | NS | NS | NS | NS | NS | NS |

<sup>a</sup>(:) represents the interaction between variables and (/) represents the nested interaction between variables.
<sup>b</sup>Significance levels for each variable are given by: \*\*\* $P < 0.001$ ; \*\* $P < 0.01$ ; \* $P < 0.05$ ; NS, $P \geq 0.05$ .

**FIG 1.**
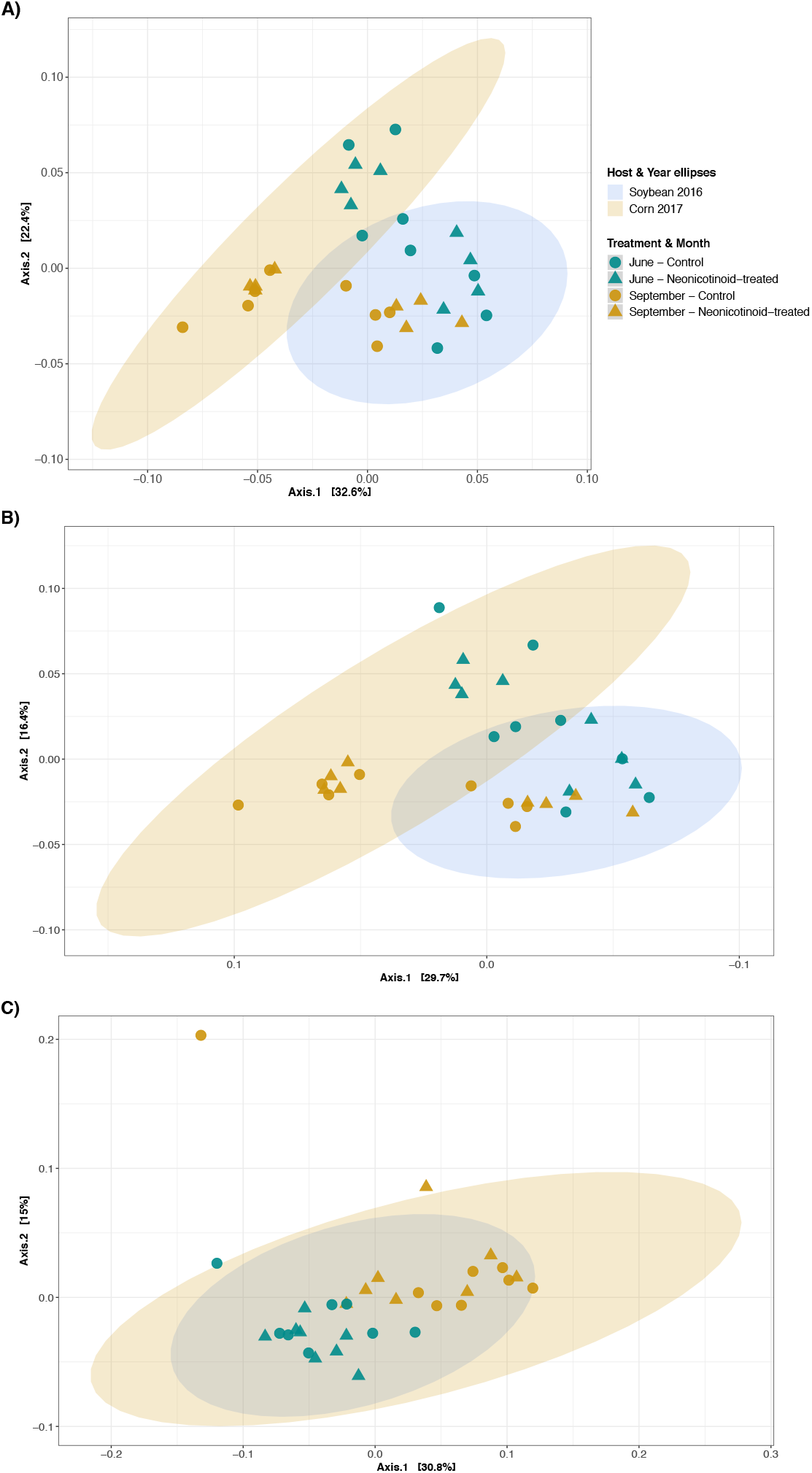
Composition variation of soil microbial expressed genes in response to neonicotinoid seed treatment and time. Principal coordinate analysis (PCoA) on Bray-Curtis dissimilarities illustrates the composition variation of soil SEED hierarchical microbial (level 4; A), RefSeq bacterial (B) and RefSeq eukaryotic (C) expressed genes between control (n = 16) and neonicotinoid-treated (n = 16) samples in a two-year soybean (2016) and corn (2017) rotation in L’Acadie, Quebec, Canada. Microbial gene expression varies among months (June: green points and September: yellow points) in control (circle) and neonicotinoid-treated (triangle) samples. Ellipses are shaded based on host species and year of cultivation (blue for 2016 soybean samples and yellow for 2017 corn samples) and represent a 99% confidence level.

Additionally, while the alpha diversity of microbial functional categories of expressed genes was not affected by year, it was significantly higher in June than September in SEED hierarchical functional categories (Shannon index mean ± SE 6.57 ± 0.02 versus 6.46 ± 0.01, Wilcoxon *P*-value < 0.0001), RefSeq bacterial functional categories (Shannon index mean ± SE 7.70 ± 0.02 versus 7.58 ± 0.01, Wilcoxon *P*-value < 0.0001) and RefSeq eukaryotic functional categories (Shannon index mean ± SE 7.14 ± 0.06 versus 6.87 ± 0.06, Wilcoxon *P*-value < 0.001).

### Effects of neonicotinoid seed treatment on differential gene expression in soil microbiome

Analysis of differential expression of genes identified no significant effect of neonicotinoid seed treatment on gene expression of all samples from both sampling times and both years of rotation together (DESeq2 adjusted *P* < 0.05). However, looking individually at each year of rotation, neonicotinoid seed treatment led to significantly increased expression of two SEED hierarchical functional categories (level 4: phycobilisome core-membrane linker polypeptide and excinuclease ABC subunit A paralog in greater Bacteroides group) in 2016, when the field was planted with soybean, and decreased expression of one SEED hierarchical functional category (level 4: inner membrane protein CreD) in 2017, in the corn field (DESeq2 adjusted *P* < 0.05, Table 3). In 2016, the expression of some RefSeq bacterial functional categories also significantly decreased (chaperone protein ClpB and heat-shock protein IbpA) or increased (protochlorophyllide oxidoreductase) in neonicotinoid-treated samples (DESeq2 adjusted *P* < 0.05, Table 3). Finally, for each sampling time, the expression of genes from a few RefSeq bacterial functional categories decreased in June (phosphonate C-P lyase system protein PhnG and beta-aspartyl-peptidase) and in September (chaperone protein ClpB) in response to neonicotinoid seed treatment (DESeq2 adjusted *P* < 0.05, Table 3).

**TABLE 3.**
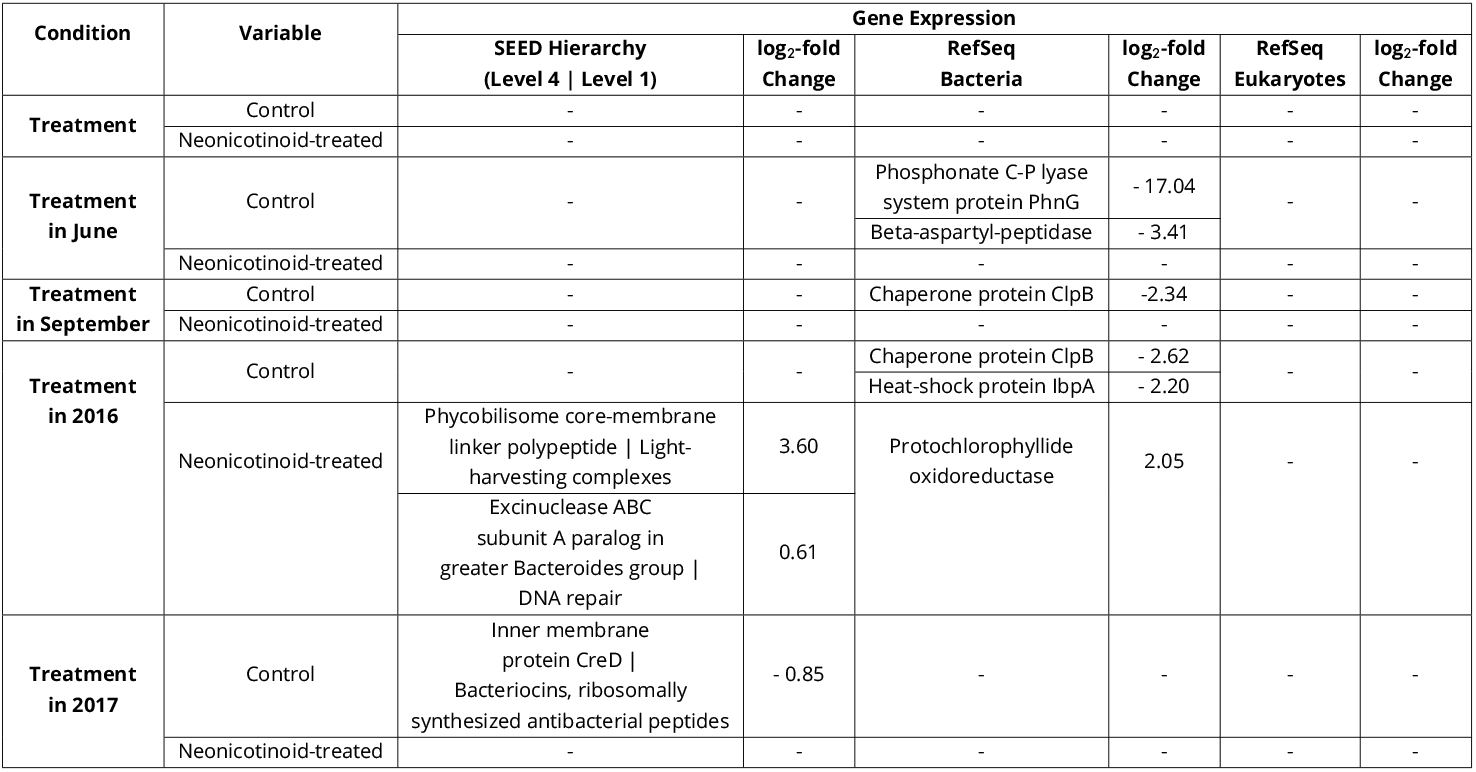
Soil SEED hierarchical microbial (level 4), RefSeq bacterial and eukaryotic expressed genes associated with control and neonicotinoid seed treatment at different times. Soil microbial genes that are significantly differentially expressed (adjusted *P* < 0.05) among different times and between control and neonicotinoid-treated samples in a two-year soybean/corn rotation in L’Acadie, Quebec, Canada identified by Differential expression analysis of sequence data (DESeq2).

While there were relatively few changes in gene expression as a result of neonicotinoid seed treatment, the expression of many soil microbial genes was impacted by time (DESeq2 adjusted *P* < 0.05). Among the SEED hierarchical functional categories (level 4), the expression of 910 genes significantly increased and 903 genes significantly decreased in 2017 versus 2016, and the expression of 516 versus 540 genes significantly increased and decreased in September versus June (DESeq2 adjusted *P* < 0.05, Tables S1Aand S1B). For example, a gene that encodes the glutathione-regulated potassium-efflux system ancillary protein KefG was significantly overexpressed in 2016 compared to 2017, as well as in September compared to June (DESeq2 adjusted *P* < 0.05, Tables S1A and S1B). Among the RefSeq bacterial functional categories, the expression of 2,250 and 2,561 genes significantly increased and decreased in 2017 versus 2016, and the expression of 1,256 versus 1,860 genes significantly increased and decreased in September versus June (DESeq2 adjusted *P* < 0.05, Tables S1C and S1D). For example, genes that encode avidin, hydroxyacid oxidoreductase and nitrogenase molybdenum-iron protein alpha chain were overexpressed in September compared to June, and also in 2016 compared to 2017 while the expression of a gene coding for pesticidal proteins significantly increased in 2017 versus 2016 (DESeq2 adjusted *P* < 0.05, Tables S1C and S1D). Finally, among the RefSeq eukaryotic functional categories, the expression of 554 and 614 genes significantly increased and decreased in 2017 versus 2016, and the expression of 322 versus 339 genes significantly increased and decreased in September versus June (DESeq2 adjusted *P* < 0.05, Tables S1E and S1F). For instance, a gene that encodes Kunitz trypsin inhibitor precursor was over-expressed in September compared to June and in 2016 compared to 2017. In addition, the expression of a gene that encodes alpha-amylase inhibitor/lipid transfer/seed storage family protein precursor increased in June versus September, and the expression of another gene encoding nematode resistance protein-like HSPRO2 increased in 2016 versus 2017 (DESeq2 adjusted *P* < 0.05, Tables S1E and S1F). Finally, based on all three microbial annotated datasets, the expression of several heat shock protein-related genes (such as heat shock protein 60, protein IbpA, chaperone protein ClpB, chaperone GroEL and chaperone GroES) increased in September, whereas the expression of the cold shock protein-related genes (such as cold shock proteins CapB, CspA and CspD) increased in June (DESeq2 adjusted *P* < 0.05, Tables S1B, S1D and S1F).

## DISCUSSION

Neonicotinoid seed treatment had weak and temporally variable effects on soil microbial gene expression in a soybean-corn agroecosystem. Conversely, time was a strong driver of the composition and diversity of soil microbial expressed genes, as expected and similar to its important effects on soil microbial taxonomic composition and diversity (44, 25). Time had a very strong effect on the expression of numerous soil microbial genes. Among them, several genes associated with cold shock protein were overexpressed in June, whereas many genes related to heat shock protein were overexpressed in September, suggesting that temporal variation in gene expression is related to changes in environmental conditions and in particularto temperature. Afew previous studies have also shown the temporal changes of soil microbial functional activities and biochemical processes in response to different agrochemical treatments, including fertilizer or pesticide application (45,46). Our results thus suggest that while gene expression in soil microbial communities is highly variable in time, these communities are either highly resistant or resilient to changes in gene expression in response to neonicotinoid seed treatment. This can be due to functional redundancy in the identity of expressed genes, despite the major variation in the taxonomic composition of these microbial communities that we have previously observed (25). Past studies have suggested that various co-occurred microbial communities may be functionally redundant. Therefore, changes in microbial taxonomic composition and diversity, especially when the community is diverse, do not necessarily affect ecosystem function (47, 48). There is thus an open question whether gene expression in soil microbial communities exhibits the pattern of functional redundancy as documented in other ecosystems (49,50,51,52, 53, 54).

Our findings indicate that the expression of some genes related to heat shock protein, metabolic processes (i.e., phosphonate break down and enzyme catalysis), and regulatory functions (i.e., respiration) decreased, while the expression of several genes related to DNA repair increased, at different time-spans in the neonicotinoid-treated samples compared to control samples. This suggests atemporally variable interaction between neonicotinoids and environmental stressors. We detected a decline in the expression of the genes related to metabolic processes, such as phosphonate C-P lyase system protein PhnG related, agene implicated in phosphonate break down, and beta-aspartyl-peptidase, which is an enzyme catalyzer, in the neonicotinoid-treated samples. This is in accordance with previous biochemical studies showing changes in soil microbial metabolic processes in response to neonicotinoid application (38,39,31,40). The observed decrease in the expression of genes such as CreD, which plays a crucial role in regulatory functions including respiration (55, 56), in the samples exposed to neonicotinoid treatment at some time points also agrees with the findings of past biochemical studies showing negative effects of neonicotinoids on soil bacterial respiration (31,57,24). Finally, an increase in the expression of genes related to DNA repair (genes encoding excinuclease ABC (subunit A)) in response to neonicotinoid seed treatment at some time points suggests that neonicotinoids may induce DNA damage in microbial cells.

Overall, despite our hypothesis that the expression of pesticide degradation-related genes would increase and the expression of nitrification-related genes decrease in response to neonicotinoid seed treatment, and previous observations of soil microbial taxonomic and physiochemical changes due to neonicotinoid application (58, 31, 40, 22, 25), we did not detect any significant shifts in the expression of genes related to biodegradation of neonicotinoids or any decline in the expression of the genes associated with nitrification. We suggest several possible explanations for this finding: First, as mentioned previously, strong temporal changes in the expression of soil microbial genes may have masked subtle effects of neonicotinoid seed treatments on gene expression. Secondly, changes in gene expression in response to neonicotinoid seed treatment may have been short-lived, and thus the gradual changes in microbial gene expression are not captured by our sampling interval. However, this seems unlikely since we sampled both early and late in the growing season. Finally, it is possible that soil microbial communities are functionally resistant or resilient, leading to few changes in gene expression in response to neonicotinoid seed treatment. Compared to measures of soil microbial community taxonomic structure (25), soil microbial gene expression seems to be less sensitive to the stress imposed by neonicotinoid application. This is probably due to the functional resilience and redundancy of microbial communities (59), and it is in line with the findings of previous studies showing less variability in microbial gene expression than taxonomic composition (49,50,52,60,54). Further validation of these findings using metabolomic analysis to quantify microbial metabolites and determine changes in microbiome metabolism in response to neonicotinoid seed treatment may help us improve our understanding of soil microbial functional dynamics and make our findings more reproducible and applicable.

Our findings are based on only two years of soybean/corn crop rotation, which makes it impossible for us to distinguish the effects of host species versus time. We did not measure environmental changes during the growing season, neither did we quantify the homogeneity of neonicotinoid concentrations across the treated samples. The changes in neonicotinoid concentration in soil overtime and among samples due to their consumption and biodegradation of neonicotinoids, the potential for an increase in the residuals of neonicotinoid and degradation products towards the end of the season and the accumulation of these products in soil over the years of rotation, and finally the changes in temperature, humidity and other environmental factors during the experience may also partially explain the effects of time on the microbial gene expression variation, and future studies will be required to distinguish among the impacts of these factors. Thus, overall we can only conclude that some combination of host species and time had important impacts on microbial communities.

The present results are based on microbial annotations against the SEED Subsystems hierarchical database and the NCBI’s RefSeq bacterial genomes and eukaryotic genomes databases. These databases are popular and reliable; however, due to a lack of standard labeling of genes, a future challenge will be to improve microbial genome databases, in particular for diverse ecosystems such as soils for which there are relatively few reference genomes and databases available and for which many gene functions remain unknown. Technological advancements such as long-read sequencing and an assembly-based approach to transcriptomics should also advance our understanding of the gene expression in large microbial eukaryotic genomes.

## CONCLUSIONS

In this study, we used metatranscriptomics of soil microbial communities to demonstrate high temporal variability but relatively weak and temporally variable effects of neonicotinoid seed treatment on soil microbial gene expression in a soybean-corn agroecosystem. In different time-spans, genes related to heat shock protein, regulatory functions (such as soil respiration) and metabolic processes (such as phosphonate breakdown and enzyme catalysis) were underexpressed in response to neonicotinoid seed treatment, whereas genes related to photosynthesis and DNA repair were overexpressed in response to neonicotinoid seed treatment. Our results demonstrate the crucial role of time and temporal changes in shaping soil microbial gene expression. To our knowledge, our study provides the first example of the impacts of neonicotinoid seed treatment on community-wide soil microbial gene expression in an experimental design representing real farming conditions. Overall, metatranscriptomic studies offer real-time and in-depth insight into the biologically active microbiomes and how microbial gene expression responds to neonicotinoid seed treatment.

## MATERIALS AND METHODS

### Study Site

The study was conducted in an experimental farm in Agriculture and Agri-Food Canada, located in L’Acadie (45°17’38.0”N; 73°C20’58.0”W), Quebec, Canada. L’Acadie is in the Canadian hardiness zone 5a and has a temperate climate and clay loam soil. In a two-year crop rotation system, we planted soybean (2016) and corn (2017) in mid-May, in 100 x 3 m plots with four replicates of non-neonicotinoid-treated (control) and neonicotinoid-treated seeds. There were four rows in each plot and the field was surrounded by two extra neonicotinoid-treated plots. All seeds were coated by three fungicides (difenoconazole, metalaxyl-M and sedaxane), in addition to 0.25 mg/seed thiamethoxam for the neonicotinoid-treated seeds. For three years before the experiment, the field had not been treated by any type of neonicotinoids and was a no-till meadow. We used glyphosate before and one month after seeding to control weeds, and in the corn field we also used 400 kg/ha NPK fertilizer (15-15-15) before seeding and 222 kg/ha N fertilizer (27.5%) one month after seeding. There were no significant differences in soil physicochemical properties (e.g., pH, etc.) across the field, nor between months or years (see more details in our previous study (25)).

### Soil Sample Collection

Each year, we retrieved 32 soil samples, two samples per plot at two sampling times (in June and September), for a total of 64 samples. For each sample, we used a sterile 2-cm diameter corer to collect soil from the upper 12-15 cm layer of the bulk soil (soil that does not adhere to plant roots) from six different spots at 10 cm around 6-10 close plants in a zigzag pattern and pooled them into one 400-500 g sample. Samples were immediately transferred to the laboratory in a cooler and kept at −80°C for RNA extraction.

### RNA extraction

We extracted RNA using the MoBio/QIAGEN RNeasy PowerSoil Total RNA Kit from 2 g of each soil sample according to the manufacturer’s instructions. To better capture the soil microbial functional variation, we extracted RNA twice from each sample and pooled them into one. We also pooled the extracted RNA of the two samples collected from the same plot (each replicate). Before and after pooling, total extracted RNA was quantified using a NanoDrop Spectrophotometer ND-1000 (NanoDrop Technologies, Inc.), and its integrity was assessed using RNA 6000 Nano LabChip Kit in microcapillary electrophoresis (Agilent 2100 Bioanalyzer, Agilent Technologies). Samples were then stored at −80^°^C until sequencing.

### Library preparation and metatranscriptomic sequencing

RNA samples with an RNA integrity number (RIN) ≥ 8.0 were sent to Genome Québec (Montreal, Quebec, Canada) for metatranscriptomic sequencing. To increase the number of sequenced mRNAs, ribosomal RNA (rRNA) was depleted from 250 ng of total RNA using Illumina Ribo-Zero rRNA Removal Kits Bacteria. Residual RNA was cleaned up using the Agencourt RNACleanTM XP Kit (Beckman Coulter) and eluted in water. The second round of ribo-depletion was done using Illumina Ribo-Zero rRNA Removal Kits (Yeast). Residual RNA was again cleaned up using the Agencourt RNACleanTM XP Kit (Beckman Coulter) and eluted in water. Complementary DNA(cDNA) synthesis was achieved with the NEBNext RNA First-Strand Synthesis and NEBNext Ultra Directional RNA Second Strand Synthesis Modules (New England BioLabs). The remaining steps of library preparation were done using the NEBNext Ultra II DNA Library Prep Kit for Illumina (New England BioLabs). Adapters and PCR primers from New England BioLabs were employed. Libraries were quantified using the Quant-iT PicoGreen dsDNA Assay Kit (Life Technologies) and the Kapa Illumina GA with Revised Primers-SYBR Fast Universal kit (Kapa Biosystems). The average fragment size (313 bp, including adapters) was determined using a LabChip GX instrument (PerkinElmer). RNA samples were finally paired-end sequenced on four lanes (eight samples per lane) on Illumina HiSeq at the Genome Québec facility (Montreal, Quebec, Canada).

### Bioinformatic analyses, quality filtering and rarefaction

We processed the metatranscriptomic data according to the standalone metatranscriptome analysis (SAMSA2) pipeline (61). We first merged the paired-end reads to make contigs using PEAR v0.9.5 (62). Then, we applied Trimmomatic v0.32 (63) (parameters: PE-phred33, SLIDINGWINDOW:4:15 and MINLEN:99) on the merged metatranscriptomes to remove adaptor contamination and low-quality sequences. Physical depletion of rRNA using the ribo-depletion kits usually eliminates about 80% of ribosomal RNA (61). To remove the rest of the rRNA, we performed a bioinformatic ribodepletion using SortMeRNAv2.0 (64). For gene annotation, we used DIAMOND aligner v2.0.4 (65, 66) to BLAST the metatranscriptomes against the SEED Subsystems hierarchical database (67) and the NCBI’s RefSeq bacterial genomes and eukaryotic genomes databases (68). We used the python scripts provided by SAMSA2 to (1) group the identified SEED genes into a four-level hierarchy of subsystems (a set of genes that are associated with each other and perform a particular biological process together), (2) aggregate the large results of annotations into summarized tables of microbial genes, and (3) calculate the metatranscriptome abundance counts for further analyses. In order to minimize the possible technical artifacts caused by the number of reads, PCR, library preparation or sequencing, we performed the following steps of data cleaning: (1) given the lack of standard labeling of genes in databases, we inspected the names of the 100 most abundant genes in each annotated dataset and gave a unique name to the same genes that were labeled differently and then combined the duplicate genes, as follows: (i) in the RefSeq-based annotations of bacteria, we replaced “DNA-directed RNA polymerase subunit beta’” with “DNA-directed RNA polymerase subunit beta”, (ii) in the RefSeq-based annotations of eukaryotes, we substituted “‘Cold-shock’ DNA-binding domain containing protein” by “cold-shock DNA-binding domain-containing protein”, and (iii) in the level 4 of SEED-based hierarchical annotations, we changed “DNA-directed RNA polymerase beta’ subunit (EC 2.7.7.6)” to “DNA-directed RNA polymerase beta subunit (EC 2.7.7.6)”; (2) then, we explored samples to verify if there are any outlier samples with a very different composition of microbial expressed genes based on Shannon diversity and the non-metric multidimensional scaling (NMDS) on Bray-Curtis dissimilarities (69); (3) we removed the rare expressed genes with fewer than five reads in the entire metatranscriptome from the RefSeq-based annotation results (respectively, 37.5% and 36.0% of the total number of bacterial and eukaryotic expressed genes); (4) we also filtered all the expressed genes annotated as hypothetical proteins (1.0% of the remaining SEED-based hierarchical expressed genes, 0.1% of the remaining RefSeq-based bacterial expressed genes, and 36.7% of the remaining RefSeq-based eukaryotic expressed genes), and (5) then we rarefied samples based on their rarefaction curves (Fig. S2) to approximately the lowest number of reads per sample in SEED-based hierarchical annotations (1,430,000 reads per sample and keeping all the samples and remaining expressed genes) and RefSeq-based annotations (1,800,000 and 260,000 reads per sample of the RefSeq-based bacterial and eukaryotic annotated datasets, respectively, which resulted in keeping all the samples and 98.5% of the remaining expressed genes). Finally, we used R to analyze these datasets.

### Statistical analyses

#### Soil SEED hierarchical microbial and RefSeq bacterial and eukaryotic functional profiling

To profile the microbial functional categories and their hierarchical levels of the soil samples collected from a two-year rotation of soybean and corn, we quantified the richness of functional categories of expressed genes (number of functional categories per sample) in SEED-based hierarchical and RefSeq-based annotated data. We also determined the ten most abundant microbial functional categories at different levels of SEED hierarchy, as well as the ten most abundant RefSeq bacterial and eukaryotic functional categories according to the total relative abundance of the annotated metatranscriptomes.

#### Effects of neonicotinoid seed treatment on the composition and diversity of soil microbial expressed genes

To study the impacts of neonicotinoid seed treatment on microbial gene expression variation, we first examined the relationships between microbial expressed genes and neonicotinoid seed treatment and time (year and month). To achieve this, we performed a permutational multivariate analysis of variance (PERMANOVA) (70) with 999 permutations on each of the annotated datasets separately using the adonis2 function of the vegan package v2.5.7 (71) in R v4.0.3 (72) (model:. ~ year/month * neonicotinoid seed treatment). We also conducted a principal coordinate analysis (PCoA) ordination based on Bray-Curtis dissimilarities on each annotated dataset to visualize the variation in microbial gene expression across the soil samples in response to neonicotinoid seed treatment. Finally, we evaluated the impacts of neonicotinoid seed treatment and time (year and month) on the alpha diversity of SEED-based hierarchical microbial expressed genes and RefSeq-based microbial expressed genes using the Shannon index. For each dataset, we examined the significance of differences in alpha diversity of expressed genes between control and neonicotinoid-treated samples using the non-parametric Wilcoxon rank-sum test (73).

#### Effects of neonicotinoid seed treatment on differential gene expression in soil microbiome

We performed a differential expression analysis of sequence data using DESeq2 (74) individually on each annotated dataset to identify microbial expressed genes that differed in abundance between all the control and neonicotinoid-treated samples, and between the control and neonicotinoid-treated samples from each sampling time during the growing season (June and September) and from each year (2016 and 2017) to study the temporal effects of neonicotinoid seed treatment on microbial gene expression, as well as between each sampling time and year regardless of the treatment to study the temporal changes of microbial gene expression. We conducted these analyses on the non-rarefied and non-normalized quality filtered and denoised data. We used the log_2_-fold changes in gene expression levels to identify genes that were differentially expressed in control versus neonicotinoid-treated samples, between months, and between years, and the Wald test with a local fit type to test the significance of the gene expression differences. Finally, we adjusted the *P*-values by applying the Benjamini-Hochberg false-discovery rate (FDR) method (75) to correct for multiple testing. We chose a significance cutoff of adjusted *P*-values < 0.05 to identify significantly differentially expressed genes between control and neonicotinoid-treated samples or across time.

### Availability of data and materials

We have deposited the raw sequences at the NCBI Sequence Read Archive (SRA accession number: PRJNA780648). Our scripts to perform the current study analyses are available in the following GitHub repository: https://github.com/memoll/metatranscriptomics.

## SUPPLEMENTAL MATERIAL

**FIG S1. Most abundant microbial functional categories.** Ten most abundant soil SEED hierarchical microbial functional categories (level 4: A), RefSeq bacterial functional categories (B), and RefSeq eukaryotic functional categories (C) in a two-year soybean/corn crop rotation in L’Acadie, Quebec, Canada. Each stack bar represents one soil sample. Mutual functional categories among the three gene profiles are represented with the same colors.

**FIG S2. Rarefaction curves of the soil microbial gene expression.** Rarefaction curves for SEED hierarchical microbial (level 4; A), RefSeq bacterial (B), and RefSeq eukaryotic (C) gene expression according to the observed number of expressed genes in soil samples of a two-year soybean/corn rotation in l’Acadie, Quebec, Canada. Each line and color represent one soil sample. The maximum sequencing coverage (x-axis: number of expressed genes) is 5,000,000 reads with cutoffs at 10,000, 50,000, 100,000, 500,000, 1,000,000 and 2,000,000 reads for SEED hierarchical microbial functional expressed genes (level 4), 10,000,000 reads with cutoffs at 50,000, 100,000, 200,000, 500,000, 1,000,000, 2,000,000 and 5,000,000 reads, and 10,000,000 reads for RefSeq bacterial functional expressed genes, and 1,500,000 reads with cutoffs at 10,000, 20,000, 50,000,100,000,200,000,500,000 and 1,000,000 reads for RefSeq eukaryotic expressed genes.

**TABLE S1.** Variation in the expression of soil microbial genes between years (2017 vs. 2016; A, C and E) and between months (September vs. June; B, D and F), based on SEED hierarchical microbial (level 4; A and B), RefSeq bacterial (C and D), and RefSeq eukaryotic (E and F) functional annotations (DESeq2, adjusted *P* < 0.05).

## ACKNOWLEDGMENTS

We are grateful to Michel Fortin, Éléonore Tremblay, Pierre-Yves Véronneau, Dave Thibouthot Ste-Croix, David Berthiaume and other colleagues at AAFC for assistance in crop cultivation and sample collection and processing. We also thank Gaston Mercier (AAFC) for performing soil physicochemical analysis. Finally, we greatly thank Golrokh Kiani (CERMO-UQAM) and Joël Lafond-Lapalme (AAFC) for their suggestions in data processing, and Geneviève Labrie (CRAM) and Annie-Ève Gagnon (AAFC) for their valuable scientific advice and suggestions.

## AUTHORS’ CONTRIBUTIONS

MP, BM, and SK conceived and designed the study; BM and SK obtained the funding; MP collected and analyzed data and wrote the manuscript; All authors critically reviewed and edited the manuscript.

